# Anxiety induced by extra-hypothalamic BDNF deficiency instigates resistance to diet-induced obesity

**DOI:** 10.1101/309930

**Authors:** Xiangyang Xie, Haili Yang, Juan Ji An, Guey-Ying Liao, Zhi-Xiang Xu, Jiwei Tan, Baoji Xu

**Affiliations:** Department of Neuroscience, The Scripps Research Institute Florida, Jupiter, FL 33458, USA; College of Animal Science and Technology, Southwest University, Chong Qing 400715, P.R. China.

**Author notes:** Correspondence: Dr. Baoji Xu, Department of Neuroscience, The Scripps Research Institute Florida, Jupiter, FL 33458, USA.

## Abstract

Anxiety disorders are associated with body weight changes in humans. However, mechanisms underlying anxiety-related weight changes remain poorly understood. Using *Emx1^Cre/+^* mice, we deleted the gene for brain-derived neurotrophic factor (BDNF) in the cortex, hippocampus, and some parts of the amygdala. The resulting mutant mice displayed elevated anxiety levels and were markedly lean when fed either chow diet or high-fat diet (HFD). The mice showed higher levels of sympathetic activity, thermogenesis and lipolysis in both brown and white adipose tissues, and higher oxygen consumption and body temperature, compared with control mice. They were still lean at thermoneurality when fed HFD, indicating elevated basal metabolism in addition to activated thermogenesis. Anxiety induced by site-specific *Bdnf* deletion similarly increased energy expenditure and minimized HFD-induced weight gain. These results reveal that anxiety can stimulate adaptive thermogenesis and basal metabolism by activating sympathetic nervous system, which enhances lipolysis and limits weight gain.

## INTRODUCTION

Anxiety is a feeling of fear and apprehension about what is to come in the absence of immediate threat, accompanied by a state of high arousal and increased vigilance (Davis et al., 2010). Occasional anxiety is believed to aid survival by increasing awareness and enabling rapid responses to possible threats (Calhoon and Tye, 2015). However, persistent and disruptive anxiety that is disproportionate to actual threat is pathological. Anxiety disorders are often linked to body weight change in humans; however, the relationship between the disorders and body weight is complex. Anxiety disorders are reportedly associated with a higher body weight in children (Anderson et al., 2006; Rofey et al., 2009), whereas some anxiety patients have been known to complain about substantial weight loss.

Anxiety is characterized by activation of the sympathetic nervous system (SNS) and the neuroendocrine system, as revealed by physiological changes such as sweating, increased heart rate, and elevated levels of corticotropin releasing factor (CRF) and glucocorticoids (Calhoon and Tye, 2015; Kreibig, 2010). As sympathetic outflow is the main determinant of thermogenesis and lipolysis in adipose tissues (Bachman et al., 2002; Harms and Seale, 2013; Rothwell and Stock, 1984), frequent or persistent sympathetic activation associated with anxiety disorders could increase energy expenditure through heightened thermogenesis and thereby reduce the risk of developing obesity. On the contrary, high levels of glucocorticoids could lead to increased visceral adiposity, as displayed in patients with Cushing’s syndrome (Charmandari et al., 2005). To date, no studies have been reported to investigate how energy balance is altered in humans or mice with elevated anxiety.

Brain-derived neurotrophic factor (BDNF) is a growth factor that plays crucial roles in neuronal development and synaptic plasticity (Huang and Reichardt, 2001; Park and Poo, 2013). Its deficiency causes anxiety-like behaviors and obesity in mice and humans (Chen et al., 2006; Gray et al., 2006; Han et al., 2008; Rios et al., 2001; Soliman et al., 2010; Xu et al., 2003). In this study, we used the *Emx1^Cre/+^* driver (Gorski et al., 2002) to abolish *Bdnf* expression in the cortex, hippocampus and some parts of the amygdala. The resulting mutant mice displayed high levels of anxiety-like behaviors, sympathetic activity, CRF expression and circulating corticosterone. Remarkably, the mutant mice were lean and resistant to diet-induced obesity (DIO) due to an increase in basal metabolism and thermogenesis in both brown and white adipose tissues. Furthermore, we found that induction of anxiety with site-specific *Bdnf* deletion in the basolateral amygdala (BLA) and surrounding area also led to similar metabolic phenotypes. These results indicate that anxiety enhances energy expenditure by stimulating thermogenesis of adipose tissues and basal metabolism through SNS activation, and consequently conveys resistance to DIO.

## RESULTS

### *Bdnf* deletion in the dorsal forebrain increases thermogenesis

To investigate potential roles of BDNF expressed outside the hypothalamus in energy balance, we sought to generate a mouse mutant, *Bdnf^lox/lox^;Emx1^Cre/+^*, by crossing floxed *Bdnf* (*Bdnf^lox/lox^)* mice (Rios et al., 2001) to *Emx1^Cre/+^* mice (Gorski et al., 2002). Using *Bdnf^klox/+^;Emx1^Cre/+^* mice in which a uniquely floxed *Bdnf* allele (*Bdnf^klox^*) expresses β-galactosidase in BDNF-expressing cells once it is recombined by Cre (Liao et al., 2012), we found that the *Emx1^Cre^* knock-in allele was able to abolish *Bdnf* gene expression in the cortex, hippocampus and some parts of the amygdala including anterior part of basolateral amygdala (BLAa), posterior part of basolateral amygdala (BLAp) and posterior part of basomedial amygdala (BMAp) (Figures 1A and S1A). We further confirmed that *Bdnf* mRNA in the cortex and hippocampus was abolished in *Bdnf^lox/lox^;Emx1^Cre/+^* mice using *in situ* hybridization (Figure S1B). When fed a chow diet, *Bdnf^lox/lox^;Emx1^Cre/+^* mice had significantly higher food intake than *Bdnf^lox/lox^* control littermates (Figure 1B), while maintaining a body weight comparable to control mice (Figure 1C). However, the *Bdnf* deletion drastically reduced fat mass without affecting lean mass (Figures 1D and S1C). Furthermore, the *Bdnf* mutant mice showed an improvement in blood glucose management as revealed by both intraperitoneal glucose tolerance test (IPGTT) and insulin tolerance test (IPITT) (Figures S1D and S1E). Given that BDNF-TrkB signaling impairment in the brain or whole body leads to severe obesity (Rios et al., 2001; Xu et al., 2003), this is a surprising metabolic phenotype from a *Bdnf* mutant.

**Figure 1.**
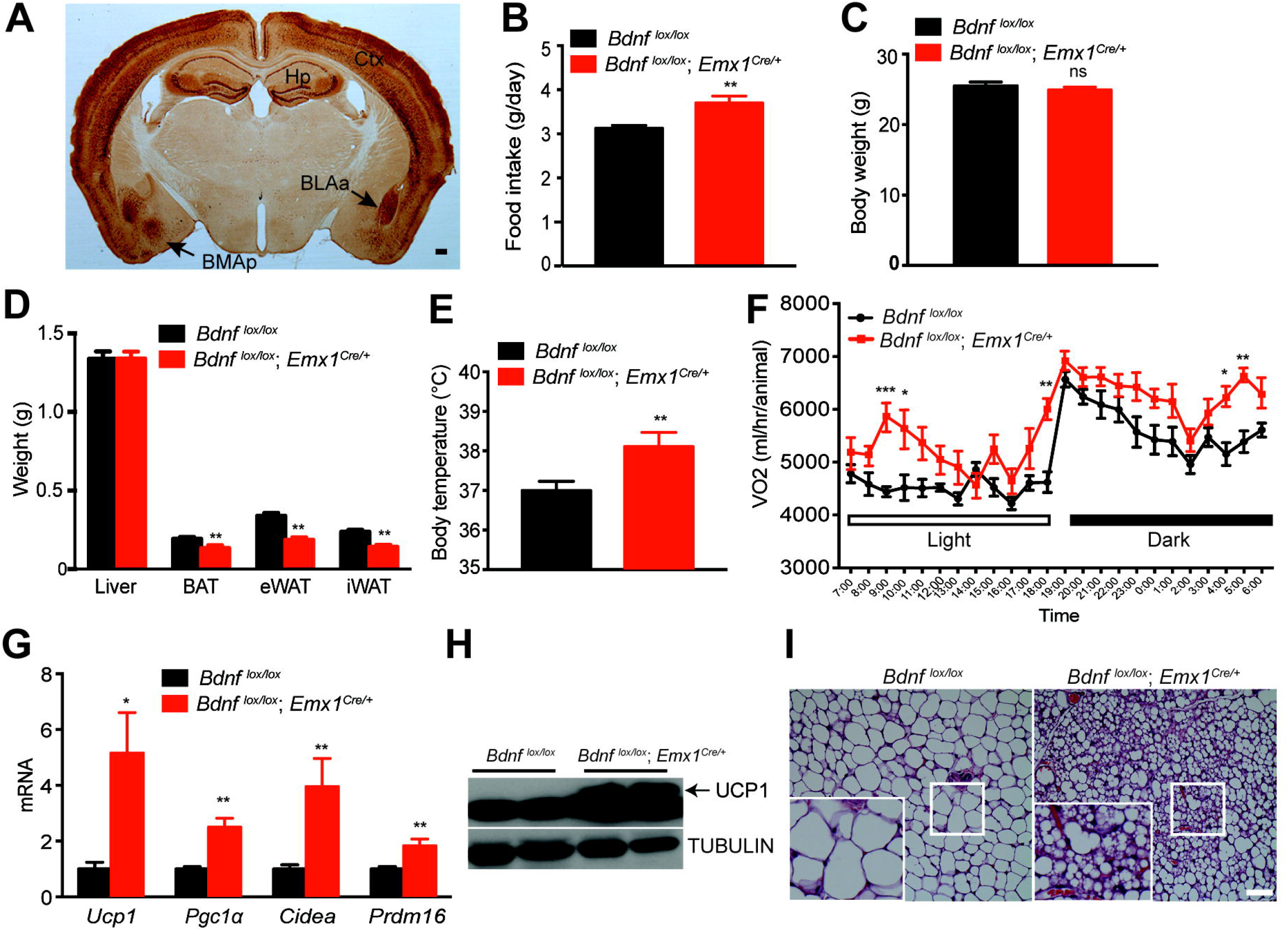
*Bdnf* deletion in Emx1-expressing neurons increased energy expenditure and induced browning in the inguinal white adipose tissue (iWAT) (A) Representative immunohistochemistry image showing expression of β-galactosidase in the brain of a *Bdnf^klox/+^;Emx1^Cre/+^* mouse. Ctx, cortex; Hp, hippocampus; BLAa: anterior part of basolateral amygdala; BMAp, posterior part of basomedial amygdala. Scale bar, 200 μm. (B) Daily intake of the chow diet. N=8-9 male mice per group; **p<0.01 by Student’s *t* test. (C) Body weight of 14-week-old male *Bdnf* mutants and control littermates raised on the chow diet. N=8-9 mice per group; ns, no significance. (D) Weight of individual tissues including the liver, brown adipose tissue (BAT), epididymal white adipose tissue (eWAT) and inguinal white adipose tissue (iWAT) dissected from 14-week-old male mice raised on the chow diet. N=8-9 mice per group; **p<0.01 by Student’s *t* test. (E) Core body temperature of mice raised on the chow diet, measured at around 4-5 pm. N=8-9 mice per group; **p<0.01 by Student’s *t* test. (F) Oxygen consumption (VO_2_) of 10-week-old male mice raised on the chow diet. N=8-9 mice per group; Two-way ANOVA followed by Bonferroni’s test: genotype, F(_1_,_336_)=99.31, p<0.0001; *p<0.05, **p<0.01 and ***p<0.001 when two genotypes were compared at individual time points. (G) Gene expression analysis of mitochondrial proteins involved in iWAT thermogenesis. N=8-9 mice per group; *p<0.05 and **p<0.01 by Student’s t test. (H) Western blotting analysis of UCP1 protein in iWAT dissected from 14-week-old male mice fed chow diet. (I) Representative images from H&E staining of iWAT dissected from 14-week-old mice raised on the chow diet. Scale bar, 20 μm. All data are presented as mean ± SEM.

Increase in food intake and reduction in fat mass in *Bdnf^lox/lox^;Emx1^Cre/+^* mice could result from increased energy expenditure. Indeed, *Bdnf^lox/lox^;Emx1^Cre/+^* mice had significantly higher body temperature (Figure 1E) and oxygen consumption during both light and dark periods (Figures 1F and S1F). The increase in energy expenditure was not a result of changes in physical activity (Figure S1G), but was in part attributable to development of brown-adipocyte-like beige cells in white adipose tissues, a process termed browning. We found that expression of genes important for thermogenesis, such as *Ucp1, Pgc1α, Cidea* and *Prdm16*, was significantly increased in the inguinal white adipose tissue (iWAT) of *Bdnf^lox/lox^;Emx1^Cre/+^* mice (Figures 1G and 1H). During thermogenesis in brown and beige adipocytes, uncoupling protein 1 (UCP1) uncouples proton gradients generated through β-oxidation of fatty acid from ATP synthesis in mitochondria and instead produces heat to maintain body temperature. Consequently, iWAT adipocytes in *Bdnf* mutant mice were smaller and contained multiple droplets (Figure 1I).

Collectively, these observations indicate that deletion of the *Bdnf* gene in the cortex, hippocampus and some parts of the amygdala induces browning of iWAT and increases energy expenditure, which leads to reduction in adiposity.

### *Bdnf^lox/lox^;Emx1^Cre/+^* mice were resistant to diet-induced obesity

Because energy expenditure was increased in *Bdnf^lox/lox^;Emx1^Cre/+^* mice, we reasoned that these animals might be resistant to diet-induced obesity (DIO). To test this possibility, we fed male mice the chow diet until 15 weeks of age and then a high-fat diet (HFD, 60% calories from fat) for additional 12 weeks. *Bdnf* mutant mice and control mice displayed similar body weights by the age of 15 weeks; however, their body weights diverged shortly after the start of HFD feeding (Figure 2A). After 12 weeks of HFD feeding, mutant mice were much lower in body weight, fat mass and lean mass than control mice (Figures 2B, S2A and S2B), although they ingested more calories than control mice (Figure 2C). Furthermore, mutant mice showed better glucose tolerance and insulin sensitivity (Figures 2D and 2E), and lower levels of liver lipid (a sign for lack of hepatic steatosis; Figures S2C and S2D) than control mice. All of these measurements show that male *Bdnf^lox/lox^;Emx1^Cre/+^* mice are resistant to HFD-induced obesity. The resistance should result from elevated energy expenditure, as the *Bdnf* deletion in Emx1-expressing neurons led to a significant increase in O_2_ consumption (Figures 2F and S2E) and core body temperature (Figure 2G). Female mutant mice also showed resistance to HFD-induced obesity, but to a lesser extent (Figures S3A-S3D).

**Figure 2.**
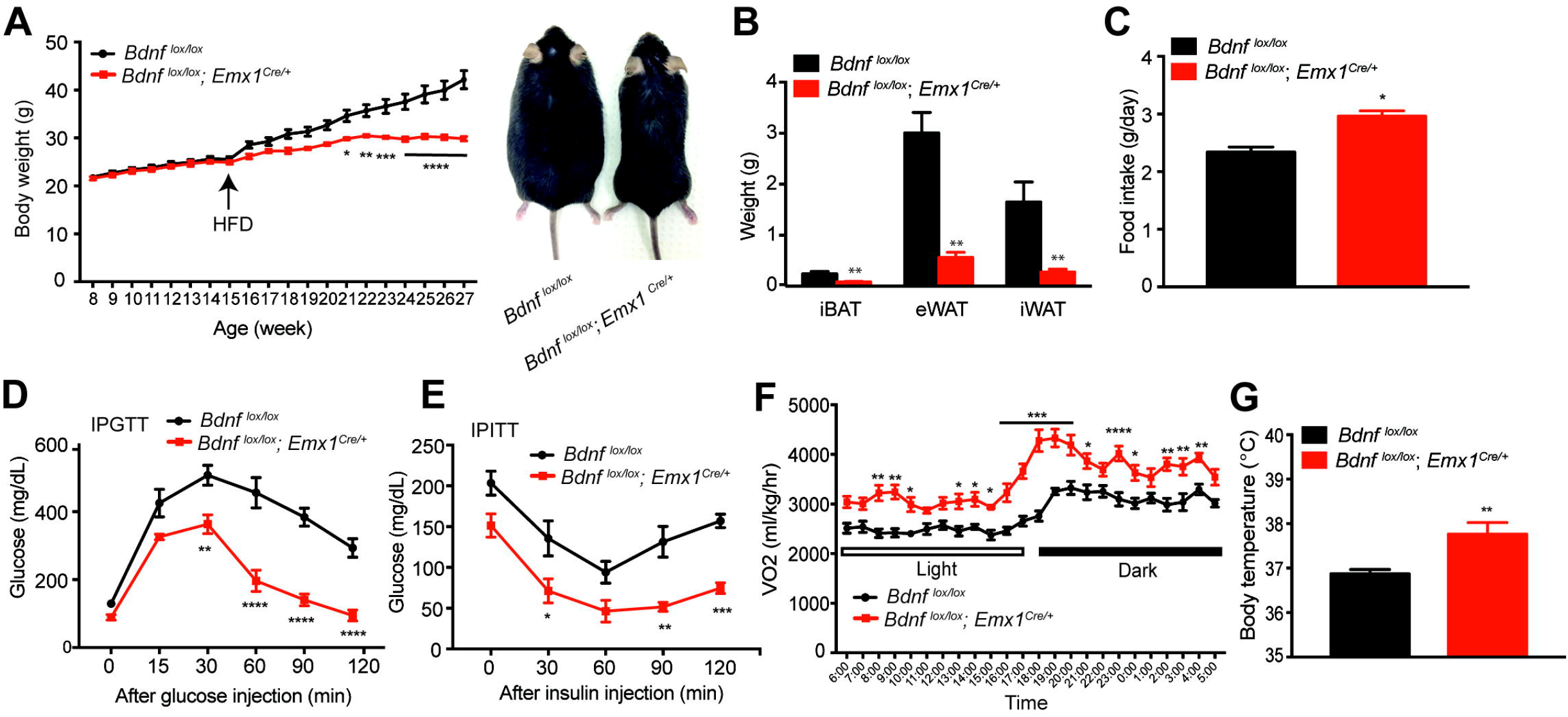
*Bdnf^lox/lox^;Emx1^Cre/+^* mice were resistant to diet-induced obesity. (A) Body weight (left) and representative pictures (right) of control and mutant mice. Animals were fed the chow diet until 15 weeks of age and then HFD for 12 weeks. N=6-9 mice per group; Two-way ANOVA followed by Bonferroni’s test: genotype, F(1,260)=139.9, p<0.0001; *p<0.05, **p<0.01, ***p<0.001 and ****p<0.0001 when comparisons were made at individual time points. (B) Weight of individual fat tissues dissected from male mice after 12 weeks of HFD feeding. N=6-9 mice per group; **p<0.01 by Student’s t test. (C) Daily intake of HFD. N=6-9 mice per group; **p<0.05 by Student’s t test. (D) Intraperitoneal glucose tolerance test (IPGTT) of male mice fed HFD for 6-7 weeks. N=6-9 mice per group; Two-way ANOVA followed by Bonferroni’s test: genotype, F_(1,48)_=116.1, p<0.0001; **p<0.01 and ****p<0.0001 when comparisons were made at individual time points. (E) Intraperitoneal insulin tolerance test (IPITT) of male mice fed HFD for 6-7 weeks. N=6-9 mice per group. Two-way ANOVA followed by Bonferroni’s test: genotype, F_(1,40)_=54.41, p<0.0001; *p<0.05, **p<0.01 and ***p<0.001 when comparisons were made at individual time points. (F) Oxygen consumption (VO2) of male mice during the first week of HFD feeding. N=6-9 mice per group; Two-way ANOVA followed by Bonferroni’s test: genotype, F_(1,312)_=372.3, p<0.0001; *p<0.05, **p<0.01, ***p<0.001 and ****p<0.0001 when comparisons were made at individual time points. (G) Core body temperature of male mice fed HFD, measured at around 4-5 pm. N=6-9 mice per group; **p<0.01 by Student’s t test. All data are presented as mean ± SEM.

### Lipid hydrolysis and thermogenesis were increased in *Bdnf^lox/lox^;Emx1^Cre/+^* mice

In consideration of thermogenesis in adipose tissues as a major component of energy expenditure in mice (Garland et al., 2011) and the observation that beige adipocytes were induced in the iWAT of *Bdnf^lox/lox^;Emx1^Cre/+^* mice fed the chow diet (Figures 1G-1I), we reasoned that increased thermogenesis in adipose tissues should significantly contribute to elevated energy expenditure in HFD-fed mutant mice. To this end, we examined morphology and gene expression in the iWAT, epididymal white adipose tissue (eWAT) and interscapular brown adipose tissue (iBAT) of HFD-fed male control and mutant mice.

In comparison with HFD-fed control mice, HFD-fed *Bdnf^lox/lox^;Emx1^Cre/+^* mice had smaller iWAT adipocytes, some of which resembled beige adipocytes with multiple lipid droplets (Figure 3A). In support of the presence of beige adipocytes, a marked increase in the protein level of UCP1 was observed in mutant iWAT (Figures 3A and 3C). Moreover, levels of mRNAs for *Ucp1* and other thermogenic genes were significantly increased in mutant iWAT (Figure 3B). Importantly, tyrosine hydroxylase (TH), the rate-limiting enzyme in the synthesis of sympathetic neurotransmitter norepinephrine, was much more abundant in mutant iWAT than control iWAT (Figure 3C), suggesting enhanced sympathetic outflow into iWAT. Consistently, the expression of β3 adrenergic receptor (ADRB3), the main receptor for norepinephrine in adipocytes, was significantly increased in mutant iWAT (Figure 3D). In accordance with increased sympathetic activity and appearance of thermogenic beige adipocytes, active and phosphorylated hormone sensitive lipase (pHSL), which is activated by adrenergic receptors through phosphorylation and is a critical enzyme in lipid hydrolysis to produce fuel for thermogenesis (Harms and Seale, 2013), was markedly increased in its level in mutant iWAT (Figure 3C). Taken together, these results indicate that *Bdnf* deletion in *Bdnf^lox/lox^;Emx1^Cre/+^* mice increases the sympathetic nerve activity (SNA) and ADRB3 activation, thereby stimulating lipolysis and adaptive thermogenesis in iWAT.

**Figure 3.**
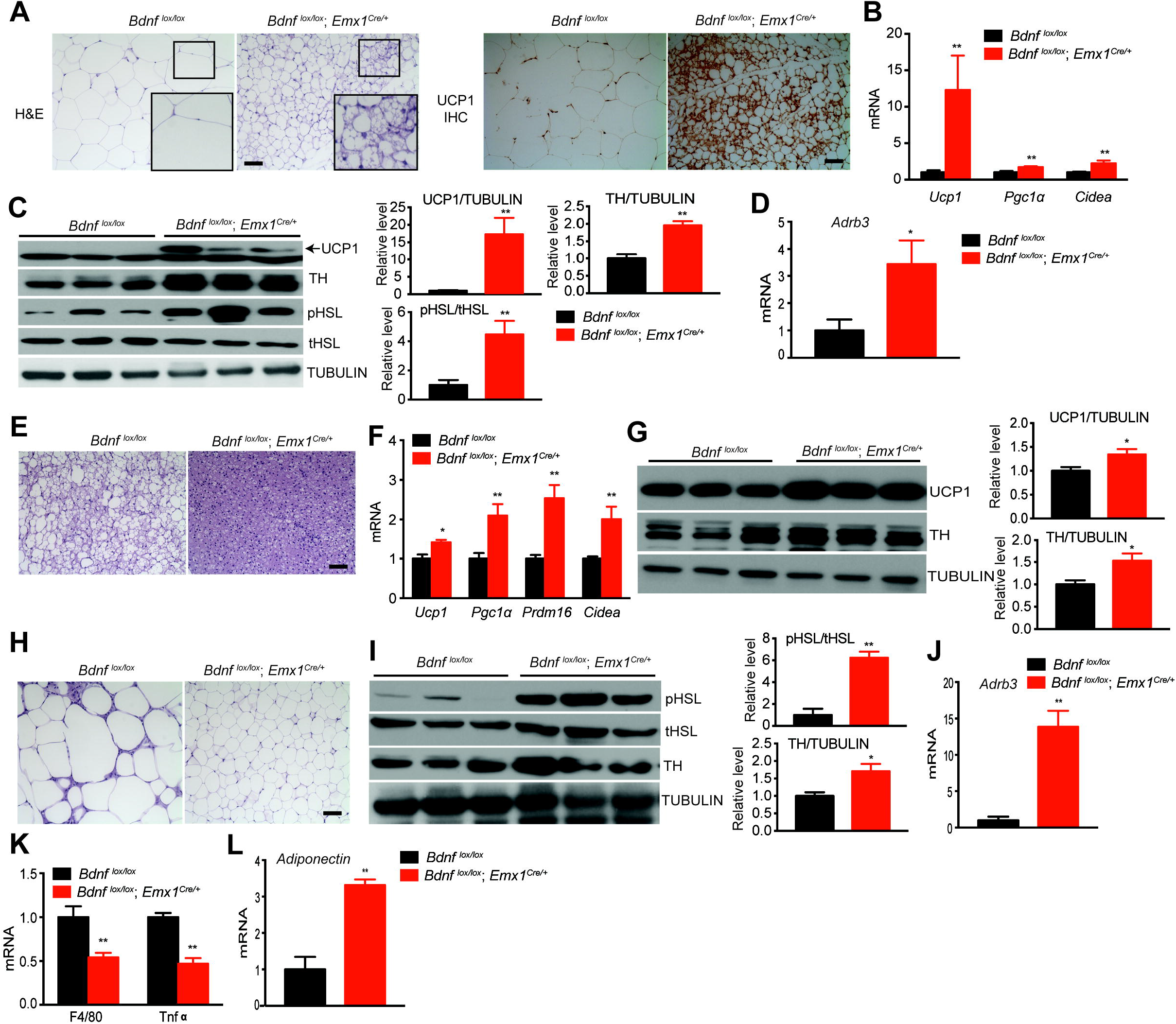
Lipolysis and thermogenesis were elevated in adipose tissues of male *Bdnf^lox/lox^;Emx1^Cre/*^* mice. All analyses were done in fat tissues dissected from mice fed HFD for 12 weeks. (A) Representative images showing H&E staining and UCP1 immunohistochemistry of iWAT. Scale bar, 20 μm. (B) qRT-PCR analysis of mRNA for thermogenic genes in iWAT. N=6-9 mice per group; **p<0.01 by Student’s t test. (C) Western blotting analysis and quantification of UCP1, tyrosine hydroxylase (TH), total hormone-sensitive lipase (tHSL) and phosphorylated hormone-sensitive lipase (pHSL) in iWAT. **p<0.01 by Student’s t test. (D) Relative levels of mRNA for β3 adrenergic receptor (*Adrb3*) in iWAT. N=6-9 mice per group; *p<0.05, by Student’s t test. (E) Representative images showing H&E staining of iBAT. Scale bar, 20 μm. (F) qRT-PCR analysis of mRNAs for genes involved in iBAT thermogenesis. N=6-9 mice per group; *p<0.05 and **p<0.01 by Student’s t test. (G) Western blotting analysis and quantification of UCP1 and TH in iBAT. (H) Representative images showing H&E staining of eWAT. Scale bar, 20 μm. (I) Western blotting analysis and quantification of HSL and TH in eWAT. *p< 0.05 and **p< 0.01 by Student’s t test. (J) Relative levels of *Adrb3* mRNA in eWAT of male mice after 12 weeks of HFD feeding. **p<0.01 by Student’s t test. (K) qRT-PCR analysis of mRNAs for proinflammatory markers in eWAT. N=6-9 mice per group. (L) Levels of adiponectin mRNA in eWAT of male mice fed HFD for 12 weeks. **p<0.01 by Student’s t test. All data were presented as mean ± SEM.

In iBAT (a key thermogenesis organ in mice (Kajimura et al., 2015)), lipid droplets were smaller in mutant mice than control mice after 12 weeks of HFD feeding, as revealed by H&E staining (Figure 3E). This morphological difference is indicative of a change in thermogenesis. In support of this view, the expression of several mitochondrial genes involved in thermogenesis, such as *Ucp1, Pgc1α, Prdm16* and *Cidea*, was significantly increased in iBAT of *Bdnf^lox/lox^;Emx1^Cre/+^* mutant mice (Figure 3F). Levels of both UCP1 and TH proteins in iBAT were also significantly increased in mutant mice compared to control mice (Figure 3G). These results indicate that elevated SNA leads to enhanced thermogenesis in iBAT of *Bdnf^lox/lox^;Emx1^Cre/+^* mice.

Like iWAT, eWAT in HFD-fed *Bdnf* mutant mice had smaller adipocytes (Figure 3H) and higher levels of TH protein, pHSL protein (Figure 3I) and *Adrb3* mRNA (Figure 3J), compared to HFD-fed control mice. These morphological and biochemical data indicate that sympathetic input and lipolysis are also elevated in eWAT of mutant mice. It is of interest that mutant mice showed a significant reduction in levels of mRNAs for F4/80 (a macrophage marker) and TNFα (an inflammatory cytokine) in eWAT, implying decreased macrophage infiltration and inflammatory response (Figure 3K). On the contrary, the level of mRNA for adiponectin was drastically elevated in mutant eWAT (Figure 3L). Given the beneficial effects of adiponectin in metabolism, this increase together with reduced inflammation in eWAT, likely contribute to a substantial improvement in insulin sensitivity observed in *Bdnf^lox/lox^;Emx1^Cre/+^* mice.

Taken together, our analyses of three fat tissues revealed that sympathetic outflow into fat tissues was enhanced in *Bdnf^lox/lox^;Emx1^Cre/+^* mice, which leads to an increase in thermogenesis of iWAT and iBAT and lipolysis, and thereby low mass of all fat tissues.

### *Bdnf^lox/lox^;Emx1^Cre/+^* mice were resistant to DIO within the thermoneutral zone

Mice are normally housed around 22°C, a temperature that is below their thermoneutrality of 30°C. As a consequence, mice live under a chronic thermal stress so that they increase heat production to compensate for heat loss (Nedergaard and Cannon, 2014). As described above, *Bdnf^lox/lox^;Emx1^Cre/+^* mutant mice displayed leanness and increased thermogenesis in adipose tissues at 22°C when they were fed either the chow diet or HFD. If elevated adaptive thermogenesis is fully responsible for leanness of *Bdnf^lox/lox^;Emx1^Cre/+^* mice, the resistance of these mutant mice to DIO should disappear within the thermoneutral zone. Otherwise, an increase in basal metabolism should contribute to the phenotype of leanness.

*Bdnf^lox/lox^;Emx1^Cre/+^* mice ingested similar amounts of calories on both the chow diet and HFD to their control littermates under a 30°C housing condition (Figure 4A). This result suggests that elevated adaptive thermogenesis leads to the observed hyperphagia in mutant mice under the 22°C housing condition. To our surprise, mutant mice were leaner than control mice even on the chow diet, and showed robust resistance to DIO (Figures 4B and S4A-S4C). Remarkably, mutant mice gained little weight even after 12 weeks of HFD exposure, whereas control mice gained a substantial weight during the same period. Consequently, compared with control mice, *Bdnf^lox/lox^;Emx1^Cre/+^* mice showed significantly improved insulin sensitivity and glycemic control (Figures 4D and 4E).

**Figure 4.**
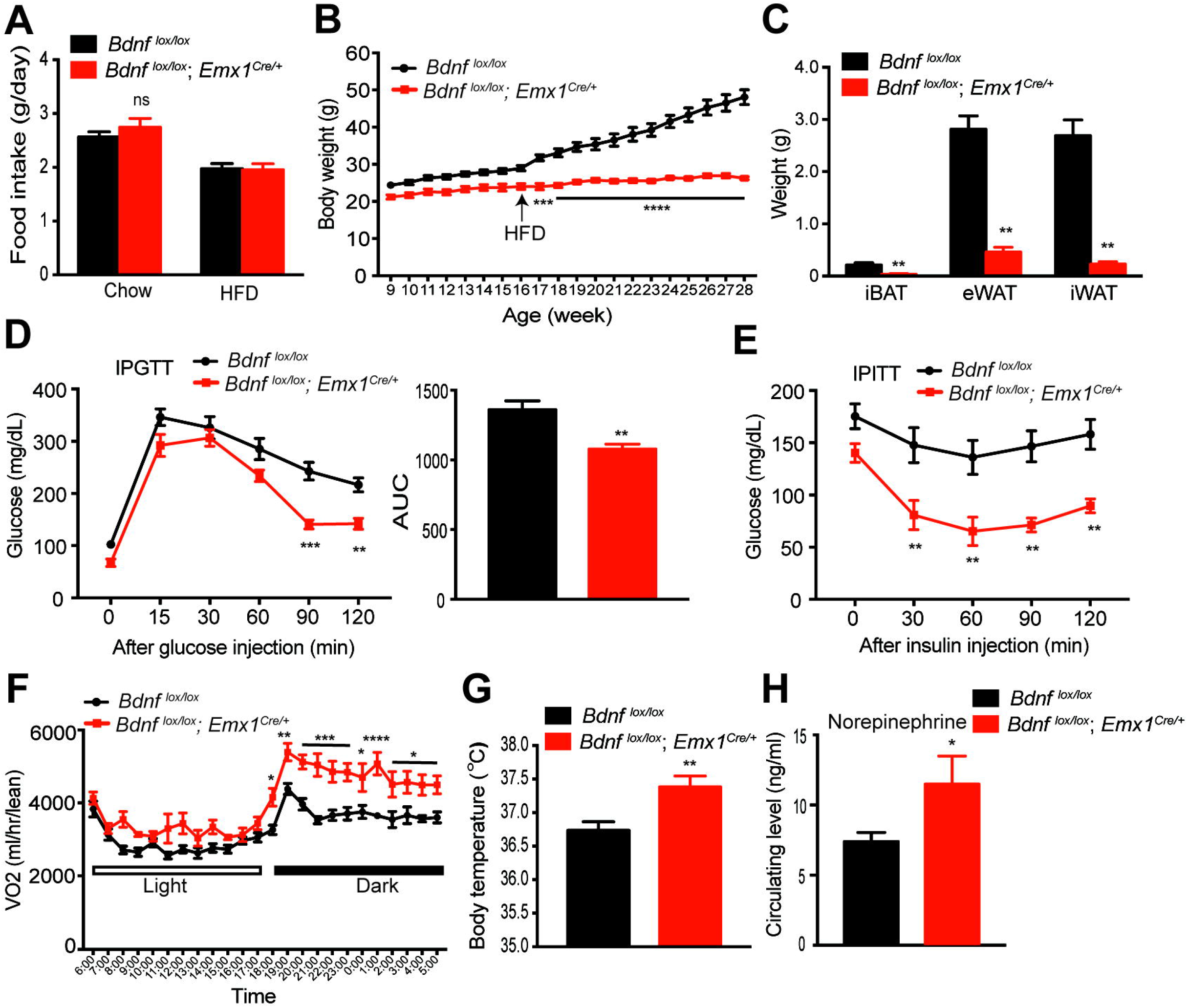
Male *Bdnf^lox/lox^;Emx1^Cre/+^* mice were resistant to DIO within the thermoneutral zone. (A) Daily intake of chow and HFD at the thermoneutral temperature (30°C). N=6-8 mice per group; ns, no significance by Student’s t test. (B) Body weight of male mice housed at thermoneutrality. Eight-week-old mice were switched to thermoneutrality from standard housing. They were maintained on the chow diet until 16 weeks of age when they started receiving HFD for 12 weeks. N=6-8 mice per group; Two-way ANOVA followed by Bonferroni’s test: genotype, F_(1,240)_=689, p<0.0001; ***p<0.001 and ****p<0.0001 when comparisons were made at individual time points. (C) Weight of individual fat tissues after HFD for 12 weeks at thermoneutrality. N=6-8 mice per group; **p<0.01 by Student’s t test. (D) IPGTT in mice fed HFD for 10-11 weeks at thermoneutrality. N=6-8 mice per group; Twoway ANOVA followed by Bonferroni’s test: genotype, F_(1,72)_=37.61, p<0.0001; **p<0.01 and ***p<0.001 when comparisons were made at individual time points. (E) IPITT in mice fed HFD for 10-11 weeks at thermoneutrality. N=6-8 mice per group; Two-way ANOVA followed by Bonferroni’s test: genotype, F_(1,51)_=52.18, p<0.0001; **p<0.01 when comparisons were made at individual time points. (F) O_2_ consumption of mice at thermoneutrality during the first week of HFD feeding. N=6-8 mice per group; Two-way ANOVA followed by Bonferroni’s test: genotype, F_(1,288)_=189.5, p<0.0001; *p<0.05, **p<0.01, ***p<0.001 and ****p<0.0001 when comparisons were made at individual time points. (G) Body temperature of mice housed at thermoneutrality. N=6-8 mice per group; **p < 0.01 by Student’s t test. (H) Concentration of norepinephrine in the plasma of HFD-fed mice. N=8-10 mice per group; *p<0.05 by Student’s t test. All data were presented as mean ± SEM.

Adaptive thermogenesis is not required to maintain core body temperature under the thermoneutral condition. Indeed, there was no difference in expression of thermogenic genes in iWAT between *Bdnf^lox/lox^;Emx1^Cre/+^* and control mice housed at 30°C (data not shown). However, mutant mice still consumed more O_2_ and had higher body temperature than control mice when they were fed HFD (Figures 4F, 4G and S4D). These results indicate that basal metabolism is also elevated in *Bdnf^lox/lox^;Emx1^Cre/+^* mutant mice. In support of this argument, circulating levels of norepinephrine, which is largely from the SNS, were increased in *Bdnf^lox/lox^;Emx1^Cre/+^* mice (Figure 4H). Therefore, leanness and resistance to DIO in *Bdnf^lox,lox^;Emx1^Cre/+^* mice result from increased sympathetic outflow to fat tissues as well as other organs, which leads to an increase in both thermogenesis and basal metabolism.

### BDNF expressed in the motor cortex did not affect energy expenditure

As described earlier, *Bdnf* expression was abolished in the cortex, hippocampus and several parts of amygdala in *Bdnf^lox/lox^;Emx1^Cre/+^* mice. We investigated the possibility that some BDNF-expressing neurons in these brain areas regulate energy expenditure through polysynaptic connection to adipose tissues. We sought to identify BDNF neurons that are polysynaptically connected to iWAT using retrograde trans-neuronal tracing. Seven days after injection of GFP-expressing retrograde pseudorabies virus PRV152 into iWAT of *Bdnf^LacZ/+^* mice, GFP-expressing neurons were found in many brain regions (data not shown). Among brain regions in which the *Bdnf* gene is deleted in *Bdnf^lox/lox^;Emx1^Cre/+^* mice, we detected many PRV152-infected neurons in layer 5 of the primary (M1) motor cortex (Figure 5A) and some infected neurons in the posterior part of basolateral amygdala (BLAp) (Figure 5D) and layer 5 of the somatosensory cortex (data not shown). Colocalization analysis of GFP with β-galactosidase indicated that many PRV152-infected neurons in the M1 cortex and BLAp were BDNF-expressing neurons (Figures 5B, 5C, 5E and 5F). Thus, some BDNF neurons in the M1 layer 5 and BLAp are polysynaptically connected to iWAT.

**Figure 5.**
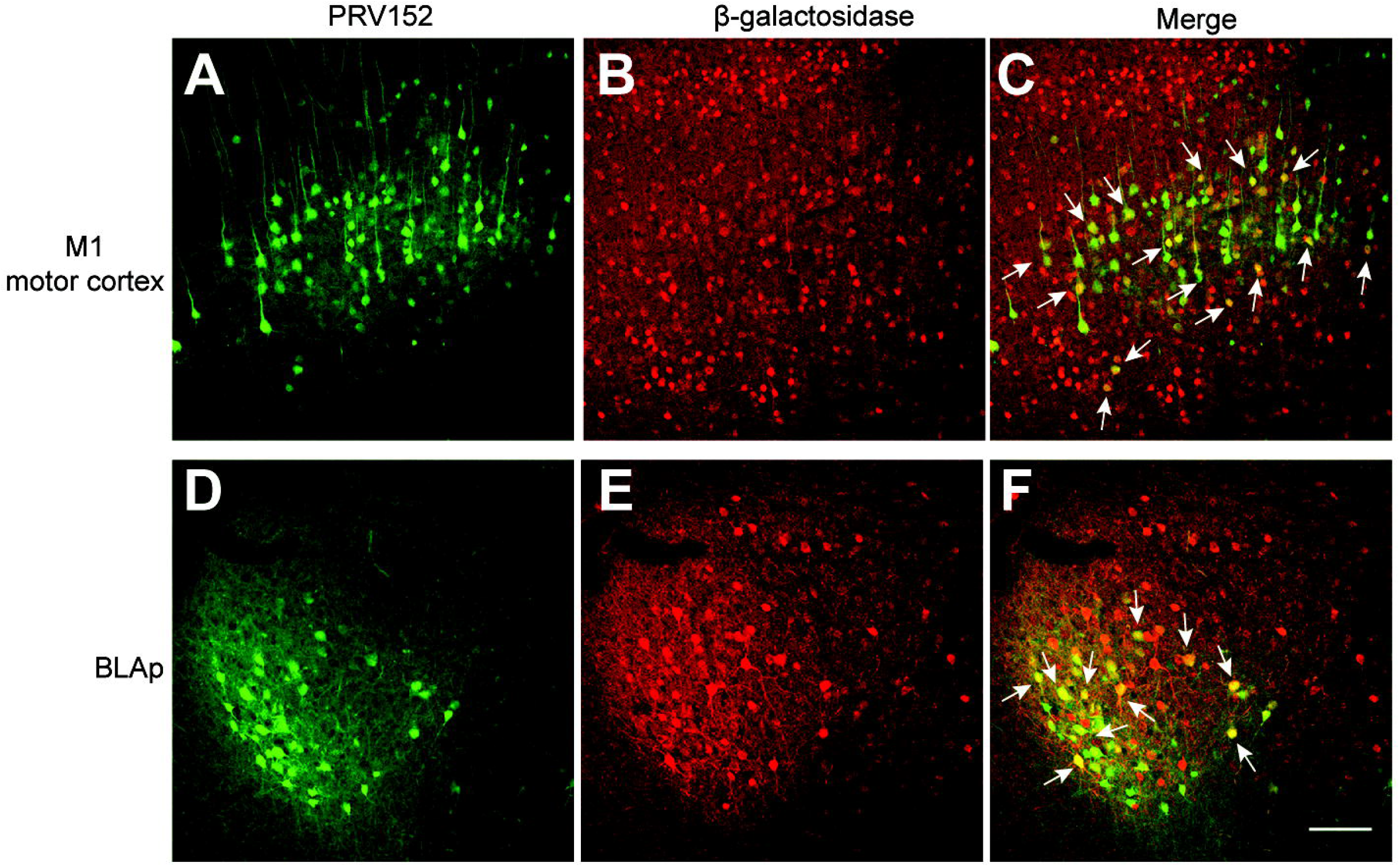
Some BDNF-expressing neurons were polysynaptically linked to iWAT. (A-C) Representative images showing infection of BDNF neurons in the M1 motor cortex of male *Bdnf^LacZ/+^* mice by GFP-expressing PRV152 injected into the iWAT. M1, primary motor cortex. (D-F) Representative images showing infection of BDNF neurons in the BLAp of male *Bdnf^LacZ/+^* mice by GFP-expressing PRV152 injected into the iWAT. BLAp, posterior part of basolateral amygdala. Immunohistochemistry against β-galactosidase revealed BDNF-expressing neurons in *Bdnf^LacZ/+^* mice. Arrows denote some neurons containing both PRV152 and β-galactosidase. Scale bar, 100 μm.

We next tested whether BDNF expressed in neurons of M1 layer 5 regulates iWAT activities and thereby accounts for iWAT browning and increased thermogenesis in *Bdnf^lox/lox^;Emx1^Cre/+^* mice. We stereotaxically injected AAV-Cre-GFP virus into M1 layer 5 of 8-week-old male *Bdnf^lox/lox^* mice to delete the *Bdnf* gene (Figure S5A). AAV-GFP virus was used as control. *Bdnf* deletion in this brain region did not have any effect on food intake, energy expenditure and body weight even after 6 weeks of HFD feeding (Figures S5B-S5D). These data suggest that ablation of BDNF expression in neurons which are connected to adipose tissues is unlikely responsible for increased energy expenditure and DIO resistance observed in *Bdnf^lox/lox^;Emx1^Cre/+^* mice.

### *Bdnf^lox/lox^;Emx1^Cre/+^* mice exhibited anxiety-like behaviors

We next investigated the possibility that abnormal behaviors could indirectly alter energy balance in *Bdnf^lox/lox^;Emx1^Cre/+^* mice. BDNF modulates synaptic transmission and plasticity (Park and Poo, 2013), and its insufficiency has been linked to cognitive impairment (Bekinschtein et al., 2008; Gray et al., 2006), anxiety-like behaviors (Chen et al., 2006; Rios et al., 2001) and aggression (Ito et al., 2011; Lyons et al., 1999). Anxiety and aggression behaviors could increase energy expenditure because they may activate the autonomic nervous system for the flight or fight response for a short period of time. Deletion of the *Bdnf* gene in the hippocampal CA3 region was found to elevate aggression (Ito et al., 2011).

*Bdnf^lox/lox^;Emx1^Cre/+^* mice were aggressive, likely due to *Bdnf* deletion in the hippocampus. We had to singly house male mutant mice in order to avoid fighting-related injuries. The *KA1-Cre* mouse line has been used to delete the *Bdnf* gene primarily in the CA3 area, resulting in an aggressive mutant (Ito et al., 2011). We regenerated this mouse mutant and detected the aggression behavior. However, male *Bdnf^lox/lox^;KA1-Cre* and control mice consumed comparable amounts of calories and gained comparable body weight after 6 weeks of HFD feeding (Figures S6A and S6B), suggesting that aggression is not the reason for low fat mass and resistance to DIO in *Bdnf^lox/lox^;Emx1^Cre/+^* mice.

We then examined whether *Bdnf^lox/lox^;Emx1^Cre/+^* mutant mice displayed anxiety-like behaviors by performing open field and light-dark exploration behavioral tests. Anxious mice incline to avoid the central zone in open field tests and the light chamber in light-dark exploration tests. In the first 5 minutes of open field tests, mutant mice traveled less distance and spent less time in the central zone than their controls (Figures 6A and 6A). *Bdnf^lox/lox^;Emx1^Cre/+^* mice also significantly decreased their entries into the central zone compared with control mice (Figure S6D), which was not a result of reduced locomotion in open field tests (Figure S6C). In light-dark exploration tests, *Bdnf^lox/lox^;Emx1^Cre/+^* mice spent less time and moved shorter distance than controls in the light chamber, and significantly reduced transition number between light and dark chambers (Figures 6C, 6D and S6E). These behavioral results indicate that anxiety is elevated in *Bdnf^lox/lox^;Emx1^Cre/+^* mice.

**Figure 6.**
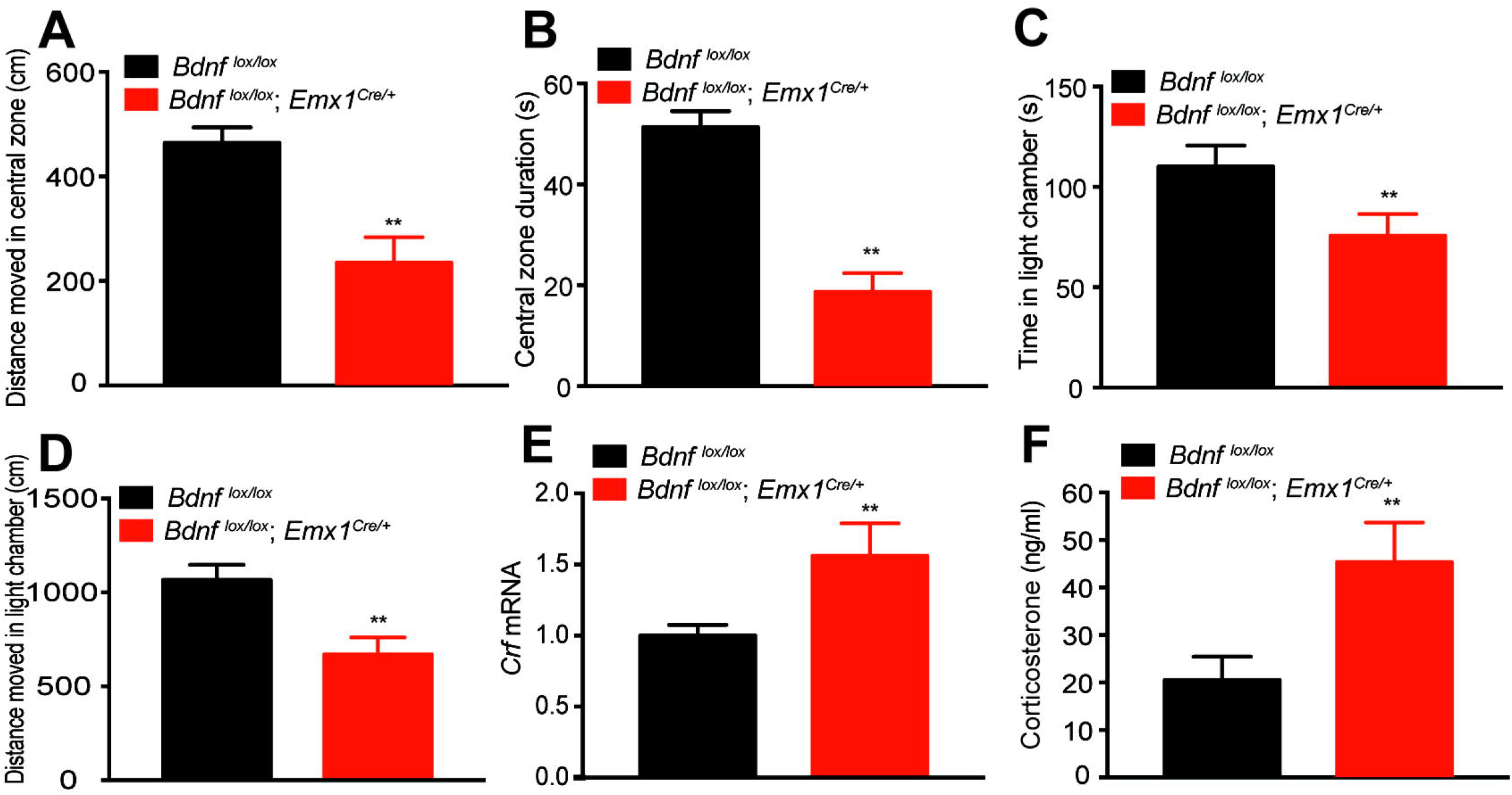
Anxiety levels were elevated in *Bdnf^lox/lox^;Emx1^Cre/+^* mice. (A and B) Distance moved in central zone and time spent in central zone (central zone duration) during the first 5 minutes of open field tests. Male mice at 10-12 weeks of age were individually housed before and during behavioral tests to avoid fighting. N=8-13 mice per group; **p<0.01 by Student’s t test. (C and D) Time spent and distance moved in the light chamber during light-dark exploration tests. N=8-13 mice per group; **p<0.01 by Student’s t test. (E) Gene expression of corticotropin-releasing factor (*Crf*) in the hypothalamus, determined by qRT-PCR. Hypothalami were dissected from mice fed HFD for 12 weeks. N=6-8 mice per group. **p<0.01 by Student’s t test. (F) Corticosterone levels in the plasma of HFD-fed male *Bdnf^lox/lox^;Emx1^Cre/+^* and *Bdnf^lox/lox^* mice. N=8-10 mice per group; **p<0.01 by Student’s t test. All data were presented as mean ± SEM.

Given a well-established connection between hypothalamic corticotropin-releasing factor (CRF) and anxiety-like behavior (Binder and Nemeroff, 2010), we examined CRF expression and production of CRF-regulated corticosterone in *Bdnf^lox/lox^;Emx1^Cre/+^* mice. Mutant mice had significantly higher levels of hypothalamic *Crf* mRNA and circulating corticosterone (Figures 6E and 6F). Hypothalamic CRF has been shown to be the central player in fibroblast growth factor 21 (FGF21)-mediated increase in the SNA and thermogenesis in adipose tissues (Owen et al., 2014).

### *Bdnf* deletion in the amygdala elevated anxiety and reduced HFD-induced weight gain

The aforementioned results indicate a strong correlation of anxious behaviors and metabolic phenotypes in *Bdnf^lox/lox^;Emx1^Cre/+^* mice. The results, however, do not rule out the possibility that the metabolic phenotypes result from an abnormal behavior we did not detect, because *Bdnf* deletion occurred in a large part of the brain in these mice and could lead to many abnormal behaviors. To address this issue, we decided to elevate anxiety by deleting *Bdnf* in a small brain region and determine whether resulting mutant mice still display resistance to DIO. We deleted the *Bdnf* gene in 8-week-old male *Bdnf^lox/lox^* mice by stereotaxically injecting AAV-Cre-GFP into the BLA (both BLAa and BLAp), which is critical for the control of anxiety-related behaviors (Calhoon and Tye, 2015). Among the mice in which the BLA was hit, we also detected AAV infection in the central nucleus of the amygdala (CeA) in the majority of them and in the medial amygdala and basomedial amygdala in a small number of them (Figure 7A). Mice injected with AAV-Cre-GFP displayed a reduced probability of entering open arms in elevated plus maze tests (Figure 7B) and decreased time spent in the light chamber in light-dark exploration tests (Figure 7C), compared with control mice injected with AAV-GFP. However, mice injected with AAV-Cre-GFP did not show abnormal behaviors in open field tests (data not shown). These behavioral observations indicate that *Bdnf* deletion in the BLA and surrounding area increases anxiety levels in mice, but not to the extent as observed in *Bdnf^lox/lox^;Emx1^Cre/+^* mice.

**Figure 7.**
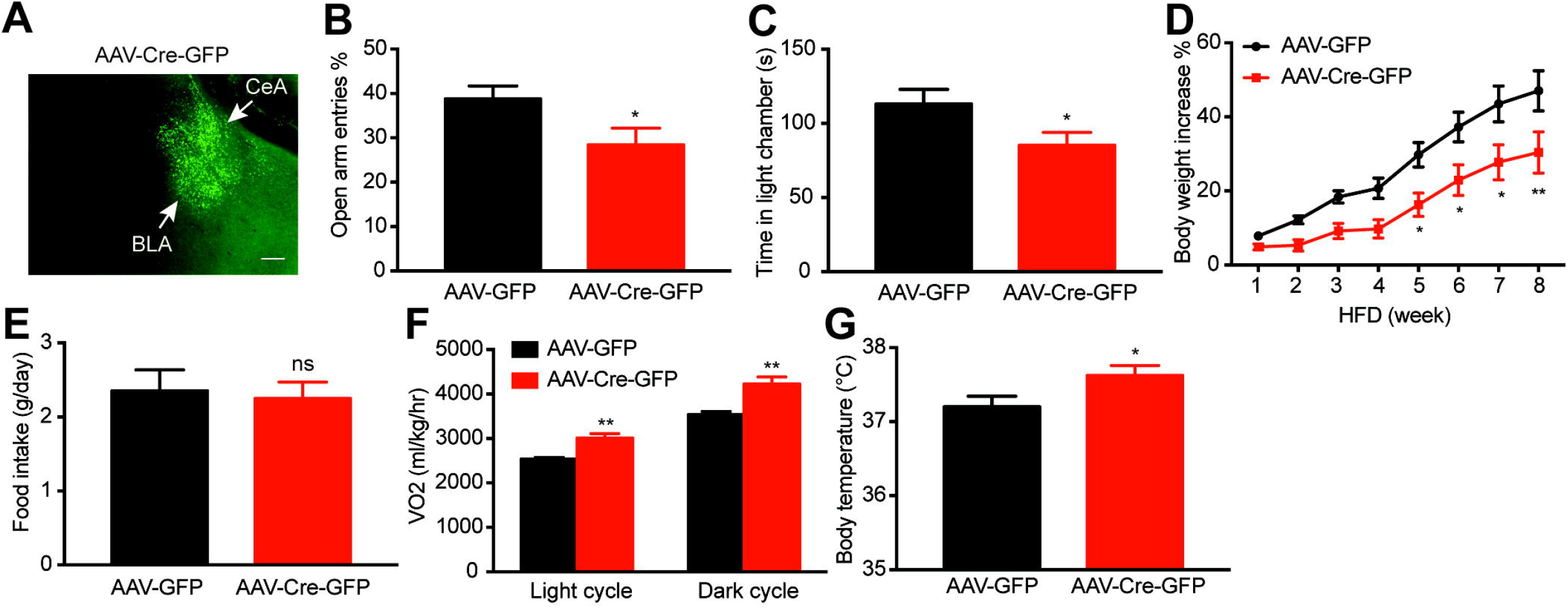
*Bdnf* deletion in the amygdala elevated anxiety and reduced HFD-induced weight gain. (A) A GFP expression image showing AAV-Cre-GFP infection in the BLA and CeA. Scale bar, 200 μm. (B) Probability of entering open arms in elevated plus maze tests. N=12-13 mice per group. *p<0.05 by Student’s t test. (C) Time spent in the light chamber during light-dark exploration tests. N=12-13 mice per group. *p<0.05 by Student’s t test. (D) Body weight increase of AAV-injected male *Bdnf^lox/lox^* mice during 8 weeks of HFD feeding. N=12-13 mice per group. Two-way ANOVA followed by Bonferroni’s test: treatment, F_(1,168)_=43.48, p<0.0001; *p<0.05 and **p<0.01 when comparisons were made at individual time points. (E) Daily HFD intake of AAV-injected male *Bdnf^lox/lox^* mice. N=12-13 mice per group. ns, no significance by Student’s t test. (F) Oxygen consumption (VO_2_) of AAV-injected male *Bdnf^lox/lox^* mice during HFD feeding. N=7-8 mice per group; **p<0.01 by Student’s t test. (G) Core body temperature of AAV-injected male *Bdnf^lox/lox^* mice during HFD feeding. N=12-13 mice per group; *p<0.05 by Student’s t test. All data were presented as mean ± SEM.

Mice injected with AAV-Cre-GFP gained smaller weight than control mice when fed HFD (Figure 7D). This was a result of increased energy expenditure, as these mice ingested normal amounts of food (Figure 7E), but had higher O_2_ consumption (Figures 7F and S7A) and core body temperature (Figure 7G) compared with control mice. However, the *Bdnf* deletion in the amygdala did not significantly alter physical activity (Figure S7B), expression of thermogenic genes *Ucp1* and *Pgc1a* in iWAT (Figure S7C), and expression of the *Adrb3* gene in iWAT (Figure S7D). Therefore, the *Bdnf* deletion in the amygdala did not alter thermogenesis in fat tissues and gave mice some resistance to DIO by increasing basal metabolism.

Collectively, we found that site-specific *Bdnf* deletion in the BLA and surrounding area of adult mice led to elevated anxiety levels, increased energy expenditure, and some resistance to DIO. This observation provides additional support for the argument that anxiety results in increased energy expenditure and resistance to HFD-induced obesity in *Bdnf^lox/lox^;Emx1^Cre/+^* mice.

## DISCUSSION

Deletion of the *Bdnf* gene in the whole brain or the hypothalamus or disruption of BDNF-TrkB signaling in the whole body leads to marked hyperphagia and severe obesity in mice (An et al., 2015; Rios et al., 2001; Xu et al., 2003). It is surprising to find that *Bdnf* deletion in the cortex, hippocampus and some parts of the amygdala led to leanness, high core body temperature and resistance to DIO in *Bdnf^lox/lox^;Emx1^Cre/+^* mice. In agreement with a report that *Bdnf* deletion in the hippocampal CA3 regions increased aggression (Ito et al., 2011), male *Bdnf^lox/lox^;Emx1^Cre/+^* mice were abnormally aggressive and constantly attacked other males when they were in the same cage. Given a critical role of BDNF in synaptic plasticity of the dorsal forebrain (Park and Poo, 2013), the mutant mice would be expected to have cognition deficits. In addition to aggression and likely cognition deficits, we found that *Bdnf^lox/lox^;Emx1^Cre/+^* mice displayed anxiety-like behaviors, indicating the importance of BDNF expressed in Emx1 neurons in the control of emotion. We generated CA3-specific *Bdnf* knockout mice as previously reported (Ito et al., 2011), and male mutant mice were aggressive but did not show leanness. Our physiological and biochemical analyses revealed that the lean phenotype in *Bdnf^lox/lox^;Emx1^Cre/+^* mice resulted from increased sympathetic activity, which elevates basal metabolism and thermogenesis in adipose tissues. There are two possible mechanisms by which the SNS is activated in *Bdnf^lox/lox^;Emx1^Cre/+^* mice: anxiety which is known to increase sympathetic activity (Calhoon and Tye, 2015; Kreibig, 2010) and inputs from Emx1-expressing neurons to the SNS through synaptic connections.

To test the possibility that some Emx1-expressing neurons inhibit metabolism of peripheral tissues through synaptic connections to the SNS and the circuit becomes dysfunctional in the absence of BDNF, we traced neurons that are connected to iWAT using pseudorabies virus. We did observe that many BDNF-expressing neurons in the M1 motor cortex were polysynaptically connected to iWAT. This polysynaptic connection has been observed before (Bartness et al., 2014; Stanley et al., 2010; Zeng et al., 2015). However, we found that deleting *Bdnf* in the M1 motor cortex had no effect on energy expenditure. Although it will be important to better understand the function of the connection between the motor cortex and iWAT in future studies, our results do not support dysfunction of this connection as the cause for the lean phenotype observed in *Bdnf^lox/lox^;Emx1^Cre/+^* mice. This exclusion approach leaves the elevated anxiety as the cause for the metabolic phenotype in these mutant mice. Our observation that site-specific *Bdnf* deletion in the BLA and surrounding area elevated anxiety and reduced HFD-induced weight gain further supports this argument. The α2 adrenergic receptor antagonist yohimbine was found to be anxiogenic in mice and rats (Figlewicz et al., 2014; Funk et al., 2008) and increase thermogenesis and lipolysis during fasting in dogs (Galitzky et al., 1991). However, these pharmacological studies did not establish a link between anxiety and metabolism; yohimbine could stimulate thermogenesis and lipolysis by acting on peripheral tissues or central neural circuits that do not control mood because α2 adrenergic receptor is widely expressed in the nervous system and non-neuronal tissues (Lorenz et al., 1990; MacDonald and Scheinin, 1995). Our genetic data provide the first evidence to indicate that anxiety increases energy expenditure and body temperature through elevated basal metabolism and thermogenesis, leading to lipolysis and resistance to DIO.

Anxiety induced by site-specific *Bdnf* deletion in the BLA and surrounding area was not as severe as the one observed in *Bdnf^lox/lox^;Emx1^Cre/^+* mice. Three reasons may account for this difference. First, site-specific *Bdnf* deletion occurred in the adult brain, while *Bdnf* deletion in *Bdnf^lox/lox^;Emx1^Cre/^+* mice happened during embryogenesis. Developmental defects could contribute to high anxiety in *Bdnf^lox/lox^;Emx1^Cre/^+* mice. Second, BDNF produced in more than the BLA and CeA is involved in mood regulation. Lastly, AAV-mediated ablation of BDNF expression in the BLA and CeA was not complete. One study reported that *Bdnf* deletion in Emx1 neurons did not elevate anxiety (Gorski et al., 2003), which could be a result of utilization of a hypomorphic floxed *Bdnf* allele. Because BDNF expression in the CeA was not altered in *Bdnf^lox/lox^;Emx1^Cre/^+* mice, we would argue that anxiety observed in mice injected with AAV-Cre-GFP results from *Bdnf* deletion in the BLA. However, interference of BDNF expression with antisense oligodeoxynucleotides in the CeA and medial amygdala, but not the BLA, was found to provoke anxiety-like behaviors in rats (Pandey et al., 2006). Therefore, further studies are warranted to define the brain structures where BDNF is produced to modulate mood. We found that both basal metabolism and thermogenesis were stimulated in *Bdnf^lox/lox^;Emx1^Cre/^+* mice, whereas *Bdnf* deletion in the BLA and surrounding area only increased basal metabolism. This observation suggests that the threshold of anxiety is higher to stimulate thermogenesis than to increase basal metabolism.

How do our findings reconcile with reports that anxiety is associated with increased body mass index in children? Whether anxiety leads to leanness or obesity could be dependent on the cause of the disorder. Several brain structures, such as bed nucleus of the stria terminalis and paraventricular hypothalamus (PVH), are involved in the control of mood and food intake (Betley et al., 2013; Calhoon and Tye, 2015; Xu and Xie, 2016). Dysfunction in some neuronal populations could lead to anxiety disorders as well as hyperphagic obesity.

Acute psychological stress transiently activates BAT thermogenesis and increases core body temperature (Oka, 2015), which involves the canonical thermoregulation neural circuit from neurons in the dorsomedial hypothalamus to the SNS via sympathetic premotor neurons in the rostral medullary raphe region (rMR) (Kataoka et al., 2014). In addition to activating BAT thermogenesis, we found that elevated anxiety also increased thermogenesis in iWAT by stimulating development of beige adipocytes in *Bdnf^lox/lox^;Emx1^Cre/+^* mice. This is somewhat similar to persistent psychological stress induced by severe burn injury, which leads to iWAT browning in both mice and humans (Porter et al., 2015; Sidossis et al., 2015). These findings suggest that anxiety-related worrying in the absence of real threats and trauma induced by severe stressors have the same impact on the SNS and adipose tissue thermogenesis. Anxiety induced by *Bdnf* deletion also increased basal metabolism; it is unknown whether psychological stress has an impact on this process.

Neural circuits of anxiety are connected to sympathetic premotor neurons in the brainstem as well as hypothalamic neurons (Calhoon and Tye, 2015). It is very likely that the canonical thermoregulation pathway is critical for anxiety-induced increase in thermogenesis and core body temperature. We found that *Crf* expression in the hypothalamus is significantly increased in *Bdnf^lox/lox^;Emx1^Cre/+^* mice. Neurons in the PVH release CRF to promote production of corticotropin in the anterior pituitary, which in turns stimulates synthesis and release of glucocorticoids in the adrenal gland. Additionally, CRF acts on neurons expressing CRF receptors throughout the CNS (Perrin and Vale, 1999). Interestingly, CRF has been found to mediate the stimulatory effect of FGF21 on energy expenditure and thermogenesis (Bookout et al., 2013; Owen et al., 2014). Therefore, the action of CRF in the brain could contribute to anxiety-induced increase in thermogenesis and energy expenditure. It would be intriguing to identify the neural circuitry through which anxiety enhances basal metabolism and thermogenesis.

## EXPERIMENTAL PROCEDURES

### Mice

All animals were on the C57BL/6 background. *Bdnf^lox/+^* (Stock No: 004339), *Emx1^Cre/^+* (Stock No: 005628) and *KA1-Cre* (Stock No: 006474) mouse strains (Gorski et al., 2002; Nakazawa et al., 2002; Rios et al., 2001) were obtained from the Jackson Laboratory (Bar harbor, ME). *Bdnf^klox/+^* and *Bdnf^LacZ/+^* mouse strains were previously described (An et al., 2008; Liao et al., 2012). Unless specified otherwise, mice were maintained at 22°C with a 12/12 hr light/dark cycle and had free access to water and food. For thermoneutral experiments, mice were housed at 30°C with a 12/12 hr light/dark cycle and had free access to water and food. The chow diet (2920x, Teklad Diets) and HFD (D12492, Research Diets) contain 16% and 60% of calories from fat, respectively. All animal procedures were approved by the Animal Care and Use Committee at Scripps Florida.

### Antibodies

Rabbit polyclonal antibody against UCP1 (ab10983) was from Abcam (Cambridge, MA). Rabbit polyclonal antibodies against total HSL (#4107) and phospho-HSL (Ser563) (#4139) were purchased from Cell Signaling (Danvers, MA). Mouse monoclonal anti-β-actin (A5441) and anti-α-tubulin (T8203) antibodies, and rabbit polyclonal anti-tyrosine hydroxylase (TH) antibody (AB152) were from Sigma (St. Louis, MO). Secondary goat anti-rabbit (HRP) (A16110) and goat anti-mouse (HRP) (#31430) antibodies were purchased from ThermoFisher Scientific (Waltham, MA).

### Immunoblotting analysis

As previously described (Mansuy-Aubert et al., 2013), animal tissues were harvested and homogenized in lysis buffer containing 1% Triton X-100, 50 mM Hepes (pH 7.4), 137 mM NaCl and proteinase inhibitor cocktail (Roche). In general, 20-30 μg of total protein was resolved in SDS/PAGE and electro-transferred to PVDF membranes. Primary antibodies were incubated with membranes at 4°C overnight. Secondary antibodies (HRP) were incubated with membranes at room temperature for 1 hr. An ECL detection system was applied to develop signals.

### Physiological measurements

Measurement of food intake, body weight, core body temperature, and fat pads was carried out as described previously (An et al., 2015; Liao et al., 2012). Body composition was assessed using a Minispec LF-50/mq 7.5 NMR Analyzer (Brucker Optics). VO2 and locomotor activity were monitored with a comprehensive lab animal monitoring system (CLAMS, Columbus Instruments). Circulating norepinephrine and corticosterone levels were measured using a Norepinephrine ELISA kit (Abnova, *#* KA1891) and a Corticosterone ELISA kit (Enzo Life Sciences, *#* ADI-900-097), respectively.

### Histology and immunohistochemistry

Adipose tissues were fixed with 4% paraformaldehyde and embedded in paraffin. H&E staining was performed on paraffin-embedded tissue sections. Liver samples were flash-frozen and sections were prepared for both H&E and Oil Red O staining. For brain immunohistochemistry, animals were anesthetized and perfused with 4% paraformaldehyde. Coronal brain sections were prepared at 40-μm thickness using a microtome. Immunohistochemistry was performed as previously described (Liao et al., 2012). The following primary antibodies were used for immunohistochemistry: rabbit polyclonal antibody against β-galactosidase (1:4000; Cappel) and rabbit polyclonal antibody against UCP1 (ab10983, Abcam).

### qRT-PCR

qRT-PCR was performed as previously described (Mansuy-Aubert et al., 2013). Briefly, tissues were homogenized in TRIzol (Invitrogen), and total RNA was extracted, reverse-transcribed to cDNA with random primers, and quantified with real-time PCR.

### IPGTT and IPITT

Intraperitoneal glucose tolerance test (IPGTT) and intraperitoneal insulin tolerance test (IPITT) were performed as described(Mansuy-Aubert et al., 2013). For IPGTT, animals were fasted overnight (16 h) and received an intraperitoneal injection of glucose (1.0 g/kg body weight). For IPITT, animals were fasted for 6 h and injected intraperitoneally with insulin (1.5 U/kg body weight). Blood was collected from the tail vain at different time points after glucose or insulin injection. A portable glucometer (OneTouch Ultra) was used to measure glucose concentrations.

#### Stereotaxic injection of AAV

AAV was stereotaxically injected into specific brain sites as described previously(An et al., 2015). In brief, mice were anesthetized with isoflurane before surgery. AAV-GFP and AAV-Cre-GFP viruses (serotype 2, UNC Vector Core) were injected bilaterally into the M1 motor cortex or BLA of 8-week-old male *Bdnf^lox/lox^* mice using a Hamilton syringe with a 33-gauge needle. The coordinate for the M1 motor cortex was AP = −0.5 mm, ML = ± 1.25 mm and DV = −1.30 mm. The coordinate for the BLA was AP = −1.5 mm, ML = ± 3.07 mm and DV = −5.15 mm. Following injections, mice received Metacam (1 mg / kg) for analgesia and were returned to their home cages.

#### Retrograde trans-neuronal tracing

Isogenic PRV-Bartha recombinant GFP-expressing PRV152 was used in the trans-neuronal tracing study as previously described(Stanley et al., 2010). Ten-week-old male *Bdnf^LacZ/+^* mice were anesthetized with isoflurane, and 5 μl of PRV152 was injected into the right inguinal white fat pad with a Hamilton syringe. The needle was left in the fat pad for 1-2 additional minutes. Seven days after PRV152 injection, mice were perfused with 4% PFA. Coronal brain sections at 40-μm thickness were stained with an antibody against β-galactosidase.

#### *In situ* hybridization

Radioactive *in situ* hybridization of brain sections was performed using ^35^S-labeled riboprobes as described before(Xu et al., 2003).

#### Anxiety-like behavioral tests

Because *Bdnf^lox/lox^;Emx1^Cre/+^* male mice always fight with other males, male *Bdnf* mutant and control mice were housed individually. Male mice were used in all behavioral tests. All behavioral tests were performed during the active phase as previously described(Chen et al., 2006; Muller et al., 2003). Mice were put into the light chamber in light-dark exploration tests. We monitored the time animals spent in each chamber, total distance mice moved in the light chamber, as well as the total number of entries into each chamber for 6 min. In open field tests, a mouse was placed at the center of the open arena and its subsequent behavior was recorded for a total of 30 min. We quantified the total distance moved in the central zone, time spent in the central zone, and total number of entries into the central zone. In elevated plus maze tests, a mouse was put at the center of the plus maze and its behavior was recorded for 5 minutes. We quantified the total distance moved, time spent at open arms, and total number of entries into open arms.

#### Statistical analyses

All data were presented as mean ± SEM. Statistical significance was analyzed using unpaired Student’s t test (Excel) or two-way ANOVA (GraphPad Prism).

## AUTHOR CONTRIBUTIONS

XX designed and performed the majority of the experiments in this study. HY and JJA started the study and made the initial observation of the lean phenotype in *Bdnf^lox/lox^;Emx1^Cre/^+* mice. GYL and JJA performed stereotaxic injection of AAV into specific brain sites. ZX and JT carried out anxiety-related behavioral tests. BX supervised the project. XX and BX wrote the manuscript.

## ACKNOWLEDGEMENTS

We thank Tingting Xia for help in tissue collection, Clint E. Kinney for help in mouse surgery, and Jessica Houtz and Shaw-wen Wu for critical comments on the manuscript. This work was supported by the grants from the National Institutes of Health to BX (R01 DK103335 and R01 DK105954). PRV152 was provided by the Center for Neuroanatomy with Neurotropic Viruses, which was supported by an NIH grant (P40 RR018604).

## COMPETING INTERESTS STATEMENT

The authors declare that they have no competing financial interests.

